# More hot air: measuring the paradox of European ecology conferences

**DOI:** 10.1101/2022.11.10.516028

**Authors:** Graham C Smith, Lineke Begeman, Alexia Coles, Emmanuelle Gilot-Fromont, Jorge R López-Olvera, Ana Vale, Barbara R Vogler, Thijs Kuiken

## Abstract

As scientists we cause an above average carbon footprint for work-related travel. International conferences are a common way for us to meet and discuss, and we do not address the environmental costs of these meetings. Knowing the costs might help us to reduce them. Here we estimated the carbon footprint of the last five physical conferences of the European Wildlife Disease Association (EWDA). We obtained the number of participants from each country for those conferences, and along with assumed travel options, commuting, accommodation, meals and printing, estimated the carbon emissions for each conference. The estimate ranged from 155 to 1205 tons CO_2_ per conference, or 0.7 to 2.4 tons per person. The outlying upper value was due to one joint global conference of the EWDA and the American mother association (WDA), which takes place every eight years. The geocentre was calculated for both the country of origin of the average conference attendants and for the society membership. The former was located in Luxembourg, and the latter near Dusseldorf, Germany, just approximately 170 km to the north. This represents the shortest total distance attendees would travel. Air transport to the conference country was the major source of CO_2_ emissions (87-97%), and the average distance flown per person increased from 1000 km (2010) to over 2000 km (2018), since later conferences were generally further from the geocentre. The biggest reduction in carbon emissions (up to 100 tons CO_2_) could therefore be achieved by decreasing air travel. This could be brought about by reducing the overall travel distance to the conference, i.e. a conference location near the membership geocentre, or – by a lesser amount – by encouraging travel to the conference by train or bus, i.e. conference location near good public travel hubs, and by promoting virtual attendance. Minor but yet substantial savings could be made elsewhere (2-6 tons CO_2_), including providing accommodation in greener hotels located closer to the conference venue and offering vegan/local food as the default option.

## 1. Introduction

It is widely acknowledged that climate change, largely caused by increasing CO_2_ emissions, is affecting the marine and terrestrial environments (e.g. Bond et al., 2019; Singh et al., 2019). Associations aimed at preserving nature and wildlife organise meetings to share knowledge and promote conservation, and thus, paradoxically adds to the increase of CO_2_ emissions due to travel, accommodation, etc. of participants at such conferences. This puts us, as members of such associations, in the awkward position of trying to understand and protect nature, while being aware of the additional damage we are inflicting due to our interests and activities. In general, we are brought up in a world where we can easily make cost-benefit analyses regarding finances. In contrast, we are not able to easily make cost-benefit analyses for environmental impact. Estimating CO_2_ emissions is an important starting point, although it is difficult to translate CO_2_ emissions to actual environmental impact. There have been calls to make conferences carbon-neutral (Bossdorf et al., 2010), and a small number of papers have started to address this issue with calls for environmental policies (Cugniere et al., 2020; Holden et al., 2017), virtual conferencing (Fraser et al., 2017; Rubinger et al., 2020) and choice of mode of travel (Fois et al., 2016). The carbon footprints of specific conferences have been calculated (e.g. Astudillo & AzariJafari, 2018; Desiere, 2016), but here we evaluate a conference series and changes over time. The objective of our study was to make future conferences of our association—the European Wildlife Disease Association (EWDA)—more sustainable by finding out which activities contributed most to the estimated CO_2_ footprint of the last five conferences. These five conferences were based at Vlieland in The Netherlands (2010), Lyon in France (2012 – a joint conference between all international sections of the Wildlife Disease Association (WDA) taking place every eight years (hereafter referred to as “global conference”), Edinburgh in Scotland (2014), Berlin in Germany (2016) and Larissa in Greece (2018).

## 2. Methods

All EWDA conferences followed the same structure, with one day allocated for student meetings and satellite meetings of sub-groups, one day for workshops and four days of conference lectures, including a half-day cultural trip. There was also a post-conference tour offered, which was not included in our calculations since it was an off-the-program optional activity. For each conference we obtained the number of participants attending from each country. We calculated the carbon emissions from these five conferences retrospectively, partly based on assumptions about people’s travel and accommodation, since more detailed information was not available.

For simplicity we assumed that the percentage of participants that travelled by air depended on their country of origin as follows: 0% for participants from the organizing country, 50% for those from a neighbouring country, 75% for those from a country bordering a neighbouring country, and 100% for those from further afield. We further assumed that local participants, and those not flying, arrived equally by train or car (assuming one person per car). For the Edinburgh conference we assumed – except for England – no immediate neighbouring country due to the insular condition of Great Britain and increased the proportion flying. We calculated all the distances from each country’s geographical centre point for simplicity, as we did not know from which airport participants flew. No account was taken of stop-over flights, or travel to the airport of origin. We assumed that people who flew subsequently travelled from the nearest international airport to the city of the conference venue by rail. We ignored the possibility that some used hire cars, as we could not retrieve any data on that.

We considered that participants stayed four nights for the conference (five for Larissa since it was not close to any airport) plus one additional night for those who attended the pre-conference workshops. We calculated daily commuting cost based on an estimated distance from recommended hotels and the means of travel most commonly used for each venue according to local organizers (e.g. walking, metro, bus or taxi). We assumed that half of the people from the host country were local and did not stay in hotels or travel (other than a similar commute to other participants). This was often the case as a high number of students attended from the local university. Travel for the within-conference trip (but not any post-conference trips) was also included, as were printing costs based on the available information on abstract production.

We calculated the carbon cost of each activity using information from various websites (Table 1). We included the information on calculating carbon costs and all data from our estimations into an Excel worksheet, so that we could easily update the total emissions. This worksheet is freely available in the supporting information and we encourage its use.

**Table 1.** The sources used in the spreadsheet to calculate the carbon footprint of transportation (per km per passenger assuming a single person per car), accommodation, meals and printing for each conference.

| Aspect | CO<sub>2</sub><br>emission<br>(kg CO<sub>2</sub> per<br>person per<br>unit) | Web site |
| --- | --- | --- |
| Travel |  | Distances travelled calculated from: |
|  |  | https://distancecalculator.globefeed.com/Distance_Between_Countries.asp |
| Flying | 0.2848/km | http://www.ghgprotocol.org/ |
| Rail | 0.01226/km | https://calculator.carbonfootprint.com/calculator.aspx?tab=4 |
| Car | 0.1368/km | https://calculator.carbonfootprint.com/calculator.aspx?tab=4 |
| Coach | 0.02801/km | https://calculator.carbonfootprint.com/calculator.aspx?tab=6 |
| Taxi | 0.15344/km | https://calculator.carbonfootprint.com/calculator.aspx?tab=6 |
| Ferry | 0.12/km | https://thewaywardpost.com/stories/carbon-footprint-of-5-travel-methods |
| Hotel | 6.5/night | http://www.epe2013.com/index.php?page=how-carbon-emission-is-computed |
| Meals | 1.67/meal | https://www.fcrn.org.uk/research-library/quantifying-carbon-footprint-catering-service-public-schools |
| Printing | 0.08/20<br>pages A4 | http://www.standardcarbon.com/2008/06/do-you-really-need-to-print-that-the-carbon-footprint-of-copy-paper/ |

Since we expected flying to be a major component of the overall carbon footprint, we further broke this down by attendance from within and outside Europe, as only the former can be markedly reduced by changing the mode of transport. For this exercise, we only considered the distance travelled: we did not take into account any variation in the carbon cost of flying due to age or make of planes.

We also calculated the geographical centre (geocentre) for EWDA members and for the average participants at each conference to examine the hypothesis that the carbon footprint of flying would increase as the conference location moved further from this geocentre. To explore this hypothesis, we took the mean attendance of Europeans from all five conferences (160 people) and then re-allocated them to each of the five conference locations to examine the difference in travel costs, using the same assumptions about means of travel for each location.

## 3. Results and Discussion

The carbon emissions from these five conferences were calculated retrospectively and various assumptions had to be made about people’s travel and accommodation. Thus, the results are estimates, but the method was comparable between conferences, and can provide enough information to suggest recommendations for future reductions of conference carbon footprint.

There was an overall trend towards higher CO_2_ emissions per EWDA conference (Figure 1) and per person (Table 2) over time. The global conference in Lyon was an outlier with more than four times higher CO_2_ emissions compared to the EWDA conferences (Figure 1). However, the joint conference meant that no North American conference took place that year.

**Figure 1.**
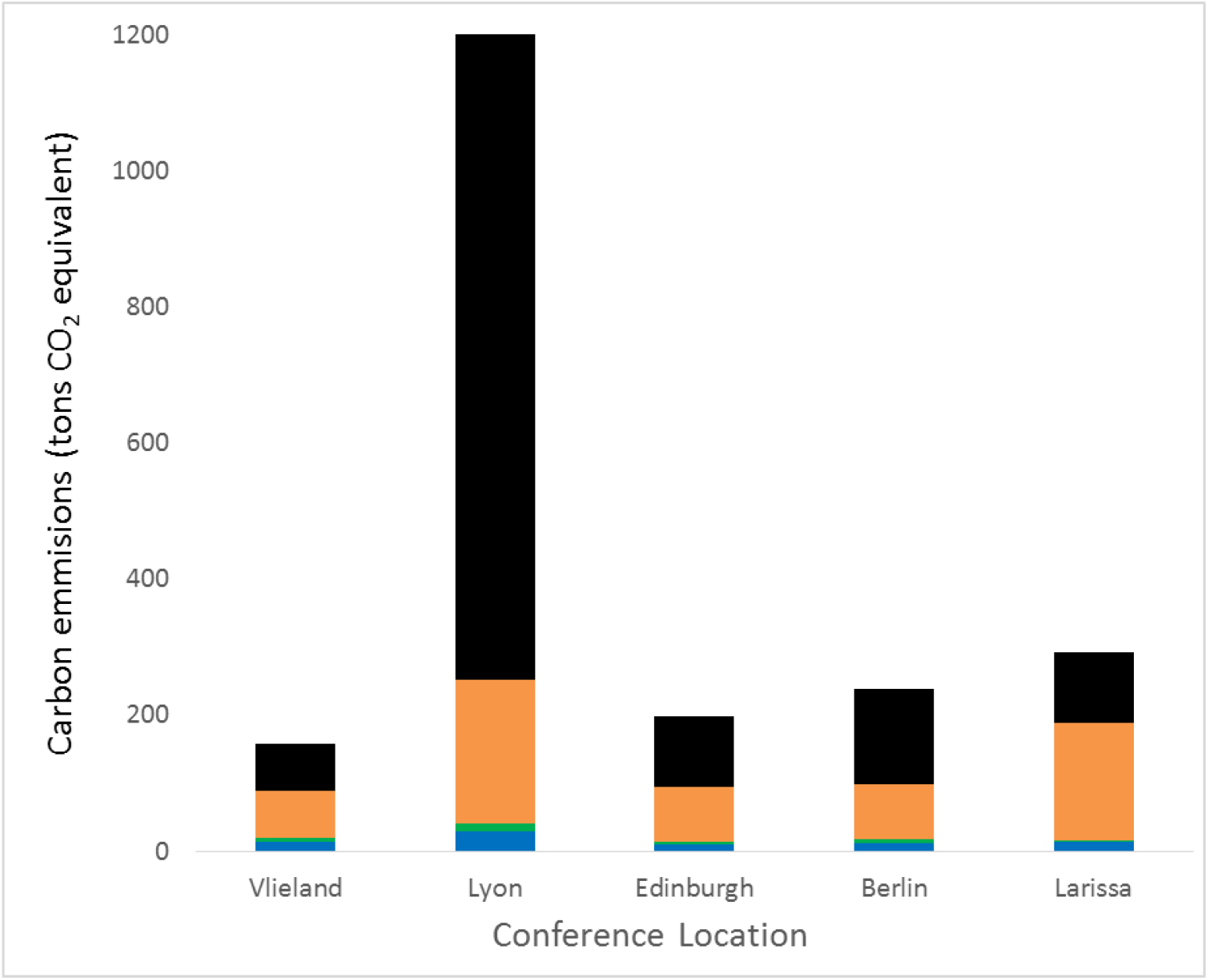
The total estimated carbon emissions (tons CO_2_ equivalent) for each conference. The top three portions show the CO_2_ emissions due to transport to the conference country (black = intercontinental flights, orange = European flights, green = other means of transportation), and the bottom portion (blue) represents all within-country costs (daily commuting, printing, accommodation, etc).

**Table 2.** Total CO_2_ costs per conference and average per person, by origin of participants.

| Year | Venue location | TOTAL (tons CO<sub>2</sub>) | Flights (tons CO<sub>2</sub>) | N total | N from country of conference | N from other European country | N from non-European country | km flown per person | tons CO<sub>2</sub> per person |
| --- | --- | --- | --- | --- | --- | --- | --- | --- | --- |
| 2010 | Vlieland | 158 | 138 | 234 | 46 | 169 | 19 | 1034 | 0.7 |
| 2012 | Lyon | 1206 | 1164 | 508 | 91 | 237 | 180 | 4023 | 2.4 |
| 2014 | Edinburgh | 197 | 183 | 198 | 80 | 97 | 21 | 1626 | 1.0 |
| 2016 | Berlin | 239 | 222 | 218 | 30 | 161 | 27 | 1789 | 1.1 |
| 2018 | Larissa | 293 | 278 | 213 | 26 | 166 | 21 | 2292 | 1.4 |

The number of participants per EWDA conference (mean 216) was very stable over time. However, the global conference attracted over 500 participants, with more participants from both inside and outside Europe (Table 2). The number of local participants attending each conference varied substantially, with both Edinburgh (80) and Lyon (91) bringing in substantially more locals than the median (46). The number of participants from outside Europe ranged from 19 and 27 for the EWDA conferences but was much higher for the global conference: 180.

The average annual CO_2_ emission per person estimated for the EU in 2017 was 7.2 tons (eurostat, 2019). Therefore, attendance at a one-week conference added from one month (conference 2010, Vlieland) to four months (global conference 2012, Lyon) of the average European’s annual carbon footprint for each attendee.

As expected, a large portion of the total carbon footprint was due to flights to the country where the conference was held (Figure 1 and Table 2). Flying accounted for between 87% and 97% of the total footprint. The other means of transport to the country of the conference (Figure 1; Table 3) was dominated by car travel. This was heavily influenced by our assumption of one person per car, so promoting methods of car sharing could lead to a noticeable reduction in the overall carbon footprint. Although train travel seemed not to be important in our scenario, we acknowledge that train transport would have had variable carbon footprint costs depending on the fuel type and source of electricity.

**Table 3.**
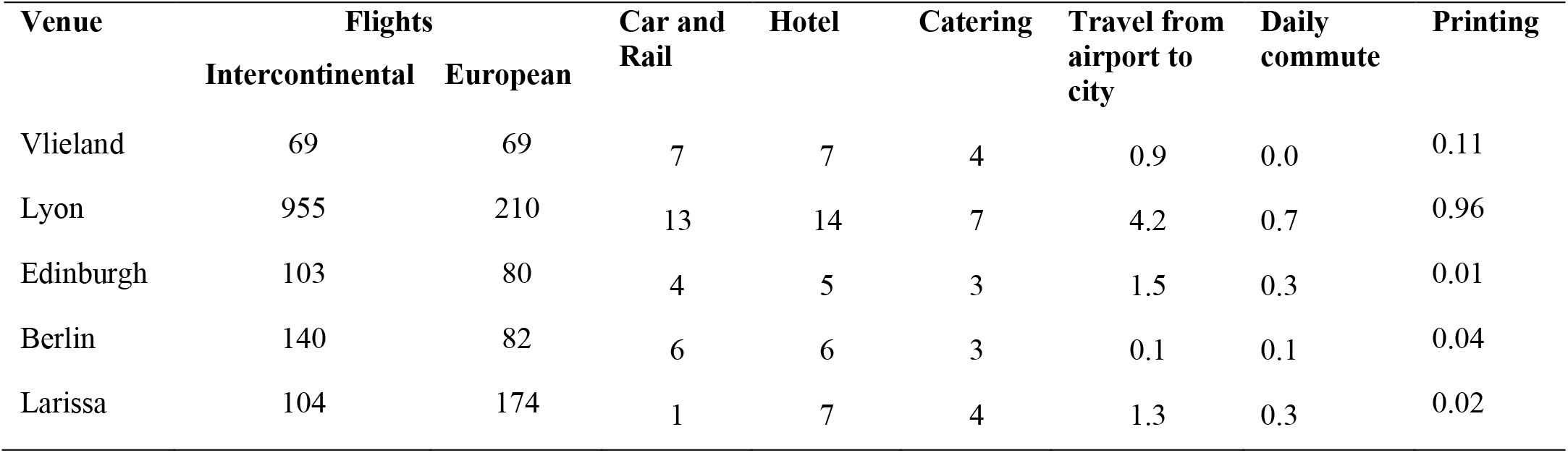
Total estimated CO_2_ emissions (tons) for the main parameters per conference. For Vlieland the daily commute was mostly by bicycle or walking, for Lyon and Berlin we assumed half used the metro, for Edinburgh all used local bus, and for Larissa all used/travelled by bus as these were supplied at each hotel.

The carbon emissions estimated for hotel accommodation, catering, the travel between airport and venue city, daily commuting, and from printing are given in Table 3.

The carbon emission resulting from the hotel accommodation is the highest remaining item (2% of total emissions on average). We used a mean value in the calculations above, but hotels with higher star ratings have a higher emissions cost per night (see link in Table 1). Nevertheless, selecting hotels with *a La Clef Verte* (Green Key) or European Ecolabel may reduce up to 50% of carbon emissions while selecting hotels with the Accor Planet21 label may reduce 5-10% of carbon emissions.

Catering accounted for 1% of total emissions. Eating vegetarian meals may reduce carbon emissions by about 32% when compared to regular meals (Cerutti et al., 2018), and this remains a substantial environmental cost for conference organisers. Encouraging vegetarian food may be helpful, for example by making vegetarian the default option and asking people to tick a box for a meat diet.

Travel to the venue city from the airport (mean 0.4% of total footprint), and daily commute costs (mean 0.06%) were lowest when the conference took place in or close to a major international travel hub (i.e. Vlieland and Berlin). This shows that locating future conferences in/near major international travel hubs can decrease CO_2_ costs. Commuting costs can also be specifically reduced by holding the conferences in well-connected small towns or locations favouring foot or bicycle commuting, as was the case in Vlieland.

The conference in Vlieland had a very low daily commute cost as it was on a small island and almost all people walked or used a bicycle. While we support the idea of conference organisers doing their best to reduce the local footprint of a conference, these data clearly indicate that even a small increase in the proportion of people choosing to take a train rather than to fly will produce a greater reduction.

The CO_2_ emissions from printing were generally negligible (mean 0.05%; Table 3), even when full paper abstracts were supplied. No reliable data on the environmental costs of the production of USB memory sticks or the use of web-based services could be obtained, so we cannot comment on whether paper abstracts, USB sticks or online abstracts are the least carbon intensive. However, paper printing may have effects on forest conservation if the conference proceedings are not on paper from sustainable sources.

The later conferences with the least number of local attendees also had the highest emissions per person, which appears to be increasing over time, when ignoring the global 2012 conference (Table 2). Lower local attendance and greater distance flown therefore combined to produce the higher carbon footprint for the later conferences (Table 2).

The geocentre calculated for the European membership was very near Düsseldorf, Germany, the geocentre for the average EWDA conference participants was in Luxembourg, just 170 km south. When examining the effect of the conference location in relation to these geocentres it was clear how much the carbon footprint increases as you move further from the geocentre, demonstrating that an ideal conference location could reduce the overall footprint by up to 100 tons CO_2_ equivalent (Table 4).

**Table 4.** The theoretical travel CO_2_ footprint, assuming the mean conference attendance by Europeans in each conference location.

| Conference | Distance from membership<br>geocentre (Düsseldorf,<br>Germany) (km) | Travel<br>footprint (tons<br>CO<sub>2</sub>) |
| --- | --- | --- |
| Vlieland | 265 | 70 |
| Berlin | 475 | 74 |
| Lyon | 617 | 134 |
| Edinburgh | 846 | 107 |
| Larissa | 1768 | 172 |

Thus, the average carbon footprint per participant has been growing over time, due to an increase in the distance to the geocentre and changes in the number of local attendees. The much greater local attendance at the Edinburgh conference accounted for the lower footprint when compared to Berlin, which was closer to the geocentre. The global conference in Lyon also attracted a greater number of European attendees than the other conferences, and this accounted for the very high European flight carbon footprint (Table 3).

## 4. Conclusions

This retrospective analysis of the carbon footprint of attending five physical EWDA conferences has revealed that the average cost could be about one ton CO_2_ per person, or even double when a large proportion of attendees come from outside Europe. The major component of this relates to flying (approximately 93%), with accommodation and catering accounting for just 2% and 1%, respectively. Traveling from airport to the conference city, daily commuting and printing costs were negligible in our case study. This strongly suggests that the biggest reduction in the total carbon footprint can be obtained by (1) locating the conference venue as close as possible to the geographical centre of the attendees to minimize the overall distance travelled by air, and (2) locating the conference venue at good travel hubs, to encourage travel by train or bus instead of by air. We expect that our calculations would be similar for many European scientific conferences leading to the same recommendations for conference organizers.

We acknowledge that, in some cases, this may be at odds with one of the conference objectives of encouraging new members, or attendance from new countries. Such considerations need to be balanced with each other. Additional consideration could be given to hosting in countries that can obtain a substantial proportion of electricity from renewable sources, and concentrating on the footprint of accommodation and catering. To reduce the carbon footprint of conferences we would further suggest the use of virtual conference rooms to deliver some talks, particularly for intercontinental attendees, and explore carbon offsetting options available. The virtual conference approach is now more common due to the covid19 pandemic, so future calculations for virtual conferences would be helpful.

## Acknowledgements

We would like to thank the organizing committees of the previous conferences for providing information and participant data on their respective conferences. All authors state that they have no conflict of interests with the content of this work and received no specific funding for this analysis. TK conceived the idea, GCS produced the spreadsheet and did the initial draft of the manuscript. All authors contributed to finding the data sources on conference attendance and footprints, contributed to the drafts and gave final approval for publication.

## Supporting Information

An Excel spreadsheet containing all the data used, which can be adapted for other conferences.

